# Tau oligomers can occur in human brains within days of a single traumatic brain injury

**DOI:** 10.1101/2025.11.18.688238

**Authors:** Shimani Kaur Bhorla, Aiste Steponenaite, Kieren Allinson, Michael Coleman, Romina Vuono

## Abstract

Traumatic brain injury (TBI) is a major risk factor for Alzheimer’s disease (AD), yet the early molecular events linking the two remain unclear. Neurofibrillary tangles (NFTs), a hallmark of AD, are observed in individuals with a history of single or repetitive head injury. Animal studies show that tau oligomers, the toxic species preceding the NFTs, appear shortly after TBI, but corresponding human data are lacking. Here, we investigated tau changes in the brains of individuals who died shortly after TBI compared to those surviving long-term. We report abnormal tau oligomers in the brain of short-term survivors whereas NFTs were the main feature in long-term survivors. These findings support a model where TBI triggers early tau oligomers formation that spread through a prion-like mechanism, seeding tau pathology and contributing to late-onset AD. The novel identification of tau oligomers in acute TBI reveals potential early disease mechanisms and therapeutic targets for AD prevention.

**Importance statement:** By revealing a potential early disease mechanism of Alzheimer’s disease (AD), this study represents a major breakthrough in our understanding, treatment and prevention of the disease. To our knowledge, this is the first report of tau oligomers in the brain of individuals suffering acute Traumatic Brain Injury (TBI). This is an important finding considering that TBI is a major risk factor of AD and tau oligomers are thought to be the most toxic species preceding tau pathology, a characteristic signature of AD. The identification of these early tau-related events after TBI provides insight into potential early AD mechanisms and offers a unique opportunity for early therapeutic intervention and prevention.

## Main

Road traffic accidents, sport injuries and falls are a significant source of Traumatic Brain Injury (TBI) affecting approximately one million people worldwide each year (**1**). Growing concern surrounds the long-term consequences of TBI with research showing that a single TBI increases the risk of developing dementia by approximately 25%, while repetitive head injuries may roughly double the risk (**2**).

Tau pathology, a characteristic hallmark of Alzheimer’s disease (AD), has been described in the brain of individuals exposed to repetitive head injuries such as boxers and National Football League players who suffered Chronic Traumatic Encephalopathy (CTE) (**3**). Furthermore, extensive tau pathology was also found in the brains of individuals surviving long-term from just a single moderate or severe TBI (**4**).

Tau is a microtubule-associated protein that promotes microtubule assembly and stability. Although the exact mechanism remains still elusive, under pathological conditions, tau becomes abnormally phosphorylated and aggregates to form insoluble intracellular deposits known as neurofibrillary tangles (NFTs) made up of abundant paired helical filaments (PHFs). Alongside the beta amyloid plaques, NFTs represent the major histopathological lesion of AD and a common feature of tau-associated neurodegenerative diseases referred to as tauopathies e.g. CTE and other conditions (**5,6**). Interestingly, the NFTs found in CTE are structurally and chemically similar to those found in AD (**7-8**).

A growing body of evidence suggests that the most toxic species may not be the NFTs, but rather pre-filamentous forms of tau, known as tau oligomers, that appear earlier in the sequence of pathological events leading to NFTs formation. Notably, tau oligomers, which may be able to enter and exit cells and propagate from disease-affected regions to unaffected areas, are thought to spread tau pathology across the brain through a prion-like mechanism involving trans-synaptic spread (**9**). In animal models of TBI, oligomeric tau species have been detected as early as 4-hours post-TBI (**10**). Moreover, tau oligomers derived from rats that underwent TBI, induced cognitive impairment when injected into brains of young, cognitively normal mice. Furthermore, consistent with a prion-like spreading mechanism, increased levels of tau oligomers were observed far from the site of injection and in multiple brain regions (**11**). However, animal studies, while extremely informative regarding the immediate effect of TBI, have limitations such as not being able to model the sheer forces causing widespread axonal injury throughout the much larger human brain. Conversely, human studies have focused on long-term TBI survivors, which do not reflect the short-term studies in animals and fail to determine whether neuro-pathologic triggers (e.g. tau oligomers) are activated at the time of injury, persist and evolve over time. To address this gap, we investigated the presence of abnormal oligomeric tau species in both, short- and long-term TBI survivors.

A total of six TBI cases were included, with survival times ranging from 24 hours to 30 years: three short-term survivors (1 to 2 days post-injury) and three long-term survivors (2 to 30 years post-injury) (**Table 1**). Multiple brain regions (**Supplementary Table 1)** were analysed by immunohistochemistry using antibodies specific for tau and beta amyloid pathology (**Supplementary Table 2**). The semi-quantitative analysis was conducted on the frontal, temporal and hippocampal regions, consistently available across all TBI cases and controls (**Supplementary Table 1**).

**Table 1.**
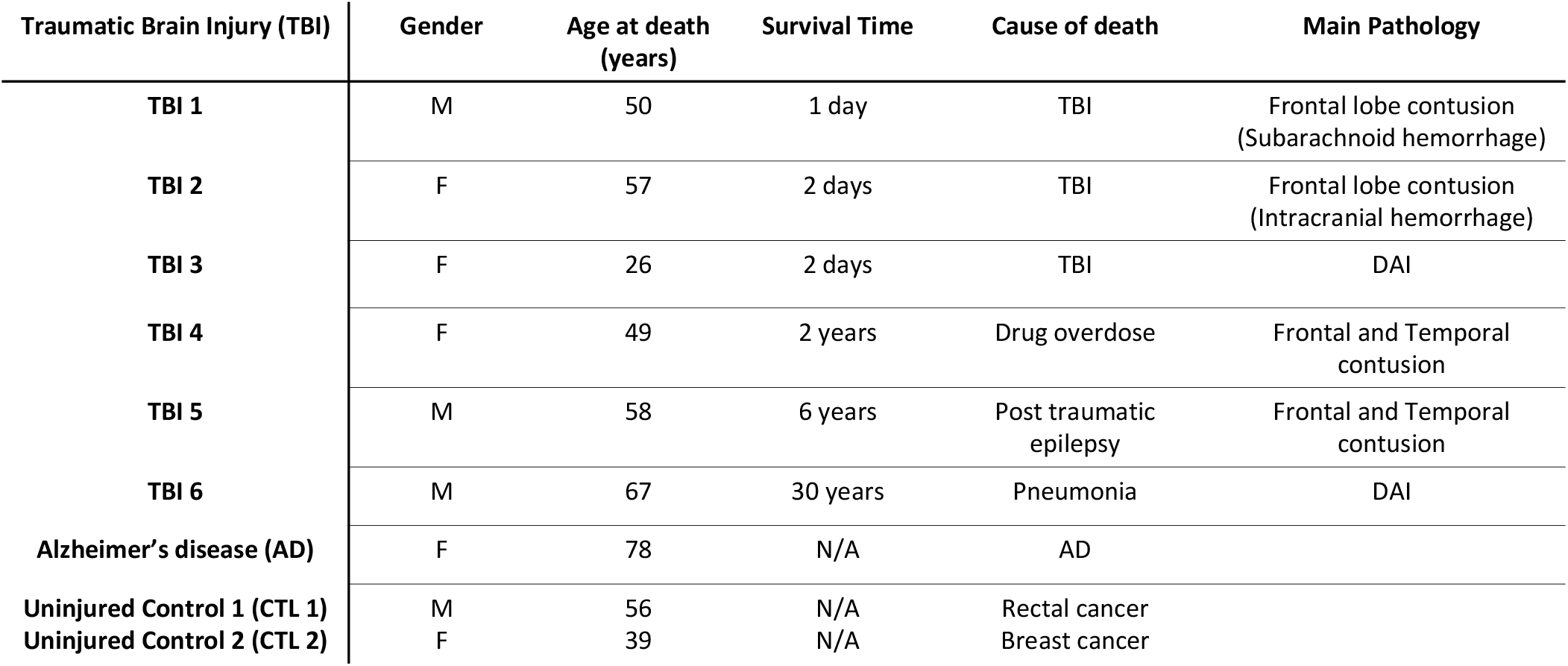
Demographic and clinical data.

We first assessed the presence of tau pathology using AT8, a monoclonal antibody that recognises abnormally phosphorylated tau aggregates and is widely used as a marker of tau-related neurodegeneration (**Supplementary Table 2**). Minimal amounts of AT8-positive neuronal deposits were detected in the brains of three short-term TBI cases (1 and 2 days) similar to observations in the uninjured controls (**Fig. 1a-c, g,h)**. Conversely, the brains of the long-term TBI cases showed increased AT8-positive staining including NFT-like aggregates and other tau inclusions (e.g. neuropil threads) similar to those seen in the AD case (**Figure 1d-f, i**). The semi-quantitative analysis confirmed that greater NFTs burden was associated with prolonged survival time (**Fig. 1j, Supplementary Fig. 1a and Supplementary Fig. 2a, Supplementary Table 3**).

**Figure 1.**
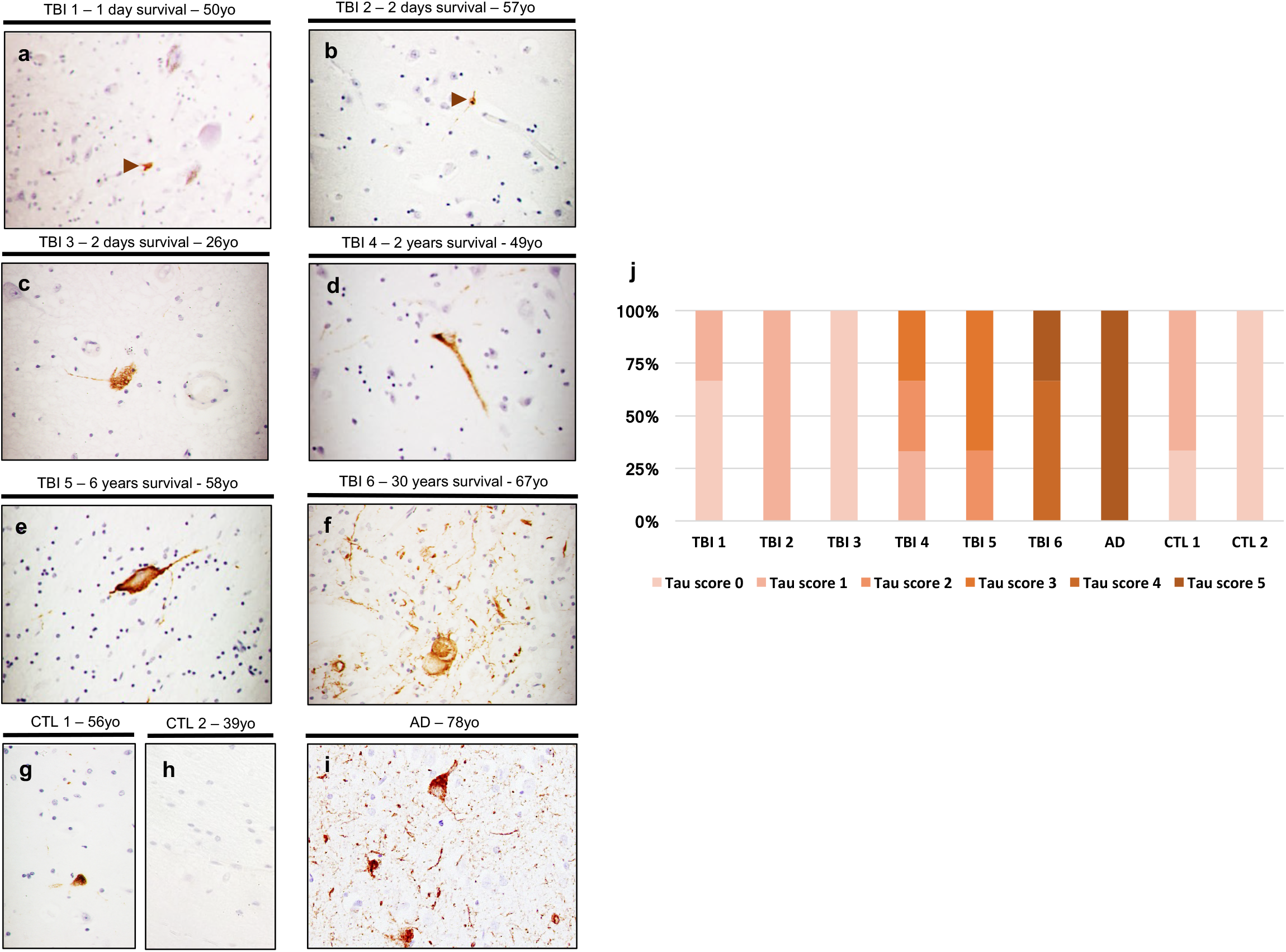
Tau pathology in the brain of short- and long-term TBI survivors and semi-quantitative analysis. Immuno-histochemical detection of pathological tau aggregates in post-mortem brain tissue from cases who have suffered a TBI (a-f) compared to AD (i) and uninjured control cases (g-h). The amount of pathological tau aggregates is minimal in the brain of short-term TBI survivors (a-c) and comparable to the uninjured controls (g-h). Conversely, long-term TBI survivors (d-f) show mature and increasing amounts of AT8 positive neuronal inclusions including NFTs, neuropil threads, and dots similarly to those observed in AD (i). (j) Semi-quantitative analysis of tau pathology determined on the three brain regions (frontal, temporal and hippocampal) available for each of the cases and categorised applying the following ranked scale: ‘nearly absent’ (score 0), ‘sparse’ (score 1), ‘moderate’ (score 2), ‘high’ (score 3), ‘very high’ (score 4) and ‘extensive’ (score 5). Results were dichotomized in either ‘nearly absent/moderate’ score (0 - 2) or high/extensive score (3 - 5) and the chi-square **g h i** test was used to assess differences between groups. Tau pathology increases with time survival since the head injury. In particular, tau pathology was ‘nearly absent/moderate’ in individuals who survived short-term whereas was ‘high/extensive’ in long-term survivors. The individual who survived 30-years (f) displayed a widespread and extensive (score 5) tau pathology similar to that observed in the AD brain (i). This fits the hypothesis that the head injury may have triggered tau changes and over time lead to the build-up and spread of pathological tau aggregates. Magnification 40X (a-i).

**Figure 2.**
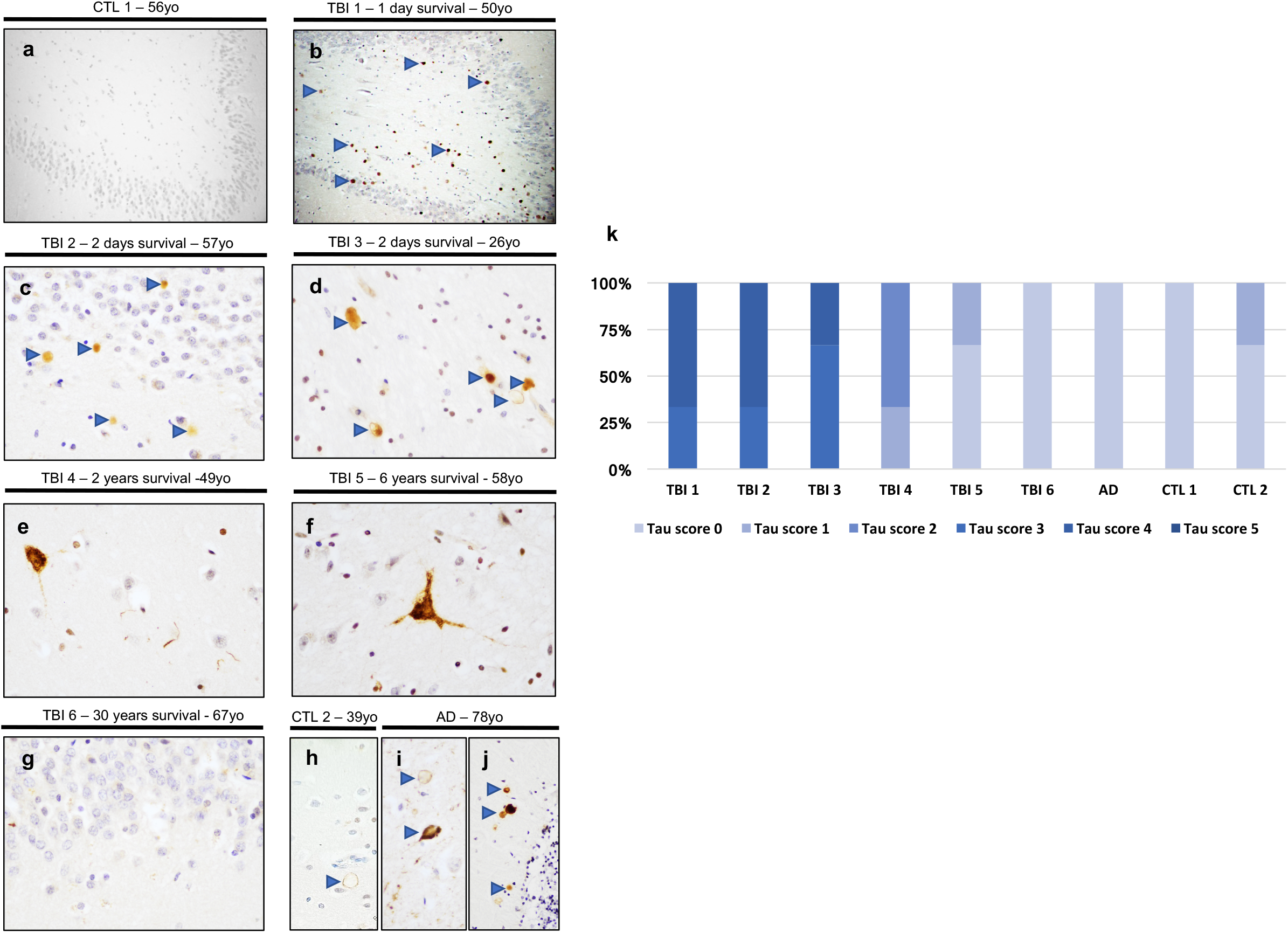
Abnormal oligomeric tau aggregates in the brain of short- and long-term TBI survivors and semi-quantitative analysis. Immuno-histochemical detection of tau oligomers in post-mortem brain tissue from cases who have suffered a TBI (b-g) compared to AD (i-j) and uninjured control cases (a,h). Oligomeric tau was detected in the brain of short-term TBI survivors (b-d) whereas was nearly absent in the uninjured control cases (a, h). The long-term TBI survival brains (e-g) didn’t show oligomeric tau but minimal amount of intermediate tau species recalling the shape of NFTs. Minimal amount of intermediate tau species and oligomeric tau was detected respectively in the AD frontal region (i) and cerebellum (j). This is in line with the tau pathology spreading which seems to affect the cerebellum later compared to other brain regions. (k) Semi-quantitative analysis of oligomeric tau was determined on the three brain regions (frontal, temporal and hippocampal) available for each of the cases and categorised applying the following ranked scale: ‘nearly absent’ (score 0), ‘sparse’ (score 1), ‘moderate’ (score 2), ‘high’ (score 3), ‘very high’ (score 4) and ‘extensive’ (score 5). Results were dichotomized in either ‘nearly absent/moderate’ score (0 - 2) or ‘high/extensive’ score (3 - 5) and the chi-square test was used to assess differences between groups. Oligomeric tau decreases as time survival increases. Tau oligomers were ‘high/extensive’ in short-term survivors and ‘nearly absent/moderate’ in long-term survivors. Of particular note, the very high amount of oligomeric tau (score 4) in the TBI short-term survivor aged 26years (TBI 3, d) compared to that nearly absent/sparse (score 0-1) of the uninjured controls (a,h), confirms that these abnormal tau species are likely to be triggered by the head injury. Magnification 20X (a,b,j) 40X (c-i).

Next, to understand what led to the tau pathology observed in the long-term TBI cases and investigate whether the NFTs were a result of a progressive build-up of oligomeric tau species triggered at the time of the head injury, we looked for the presence of tau oligomers in short- and long-term TBI survivors. Using the T22 antibody (**Supplementary Table 2**), originally characterised by *Lasagna-Reeves et al* (**12**) and widely used to detect tau oligomers in human tissue (**13-14**), a distinct granular staining was found across multiple brain regions in short-term TBI cases (**Fig. 2b-d**). This T22-positive staining was nearly absent in comparable regions of the age-matched uninjured control (**Fig. 2a**). Notably, the very high amount of tau oligomers in the brain of the short-term TBI case aged 26 (**Fig. 2d**), almost absent in the uninjured control brains aged 39 and 56 (**Fig. 2h,a**), indicates that these aggregates can occur independently of age-related processes.

In contrast, the brains from the three long-term TBI cases (2, 6 and 30 years) did not display oligomeric tau species but sparse T22 positive aggregates, most likely to be an intermediate oligomeric tau stage preceding the NFTs and still detectable by the T22 antibody (**Fig. 2e-f**). Similar intermediate tau species were occasionally detected also in the AD brain (**Fig. 2i**) as well as tau oligomers which were restricted only to the cerebellum (**Fig. 2j**). This supports a model in which NFTs arise from earlier-stage tau oligomers that may propagate via a prion-like mechanism with the cerebellum being the latest brain region showing tau oligomers. This may explain why the cerebellum is relatively spared by tau pathology and AD. The semi-quantitative analysis showed a decline in tau oligomers burden with increasing survival time post-injury (**Fig. 2k, Supplementary Fig. 1b and Supplementary Fig. 2b, Supplementary Table 3**). This pattern would be consistent with a model in which oligomers form acutely after TBI and gradually mature into NFTs-like aggregates over time.

Finally, we investigated the presence of beta amyloid plaques, the other main pathological hallmark of AD, using the antibody MOAB-2 (**Supplementary Table 2**) which specifically detects beta amyloid pathology. As expected, beta amyloid pathology was null in the uninjured age-matched control brain and widespread in AD (**Supplementary Fig. 3a-b**). In contrast, beta amyloid pathology was inconsistently observed across the TBI cohort and was detected only in the frontal brain region of the individual aged 50 and with shortest survival time (TBI 1, 1-day survival) (**Supplementary Fig. 3c-d**). This finding is consistent with previous studies reporting that TBI-associated plaques can develop within hours post-injury and may resolve over time (**15**) whilst plaques in AD develop slowly and are predominantly found in the elderly. However, it is noteworthy that beta amyloid pathology was absent in the other short-term TBI cases (TBI 2 and 3, 2-day survival), despite TBI 2 being of similar age (57 years) and having a similar injury (frontal contusion) to TBI 1 (**Table 1**). One possible explanation might lay in genetic factors playing a role in TBI associated plaques formation. A larger cohort of short- and long-term TBI cases is crucial to clarify the temporal dynamics of beta amyloid pathology following TBI, and to determine whether its appearance is mechanistically linked to tau pathology and/or influenced by individual genetic factors.

In summary, despite the limitation of a small sample size and availability of formalin-fixed paraffin-embedded tissue only, this study provides the first evidence in humans of abnormal early tau-related events after TBI and how they may evolve over time. Notably, tau oligomers can appear within hours of a single TBI, mirroring observations in animal models and regardless of age (**Fig. 2, Supplementary Fig. 1 and Supplementary Fig. 2**) (**11**). Conversely, the presence of oligomeric tau was minimal in the brains of those who survived for years after TBI (2- to 30-years), whose brains instead showed increasing numbers of NFTs (**Fig. 1, Supplementary Fig. 1 and 2**).

These findings are consistent with recent cell-culture studies showing that small globular tau aggregates form within hours after seeding but decrease at later stages, as mature fibrillary tau assemblies develop (**16**).

We therefore hypothesise that abnormal oligomeric tau species form acutely after head injury, spread in a prion-like fashion, and seed the AD-like tau pathology seen decades later. Indeed, the high burden of oligomeric tau found in the short-term TBI case aged 26 (**Fig. 2**) lends support to this hypothesis. On the other hand, the NFTs seen in the long-term TBI cases, and in particular the AD-like tau pathology observed in the individual who survived 30 years (**Fig. 1**), fits the model of a widespread tau pathology as the result of a build-up and spread of tau oligomers triggered by the head injury three decades earlier. These findings are further supported by emerging evidence from human dementia studies showing tau pathology spread across brain regions (**13-14**). Indeed, the possible sequence of events triggering the abnormal formation of tau oligomers may start soon after the head injury. In particular, the rapid acceleration–deceleration forces from the trauma can stretch and tear brain components, particularly axons, which are highly vulnerable due to their length and structural fragility. Axonal swelling and disconnection lead to diffuse axonal injury responsible for cytoskeletal changes and dysfunction (**17**). Indeed, ultrastructural analysis of axons shows immediate breakage and buckling of microtubules post-injury, which triggers progressive microtubule network disassembly (**18**). This in turn may cause tau detachment from the microtubules, imbalance between kinases and phosphatases and consequent hyper-phosphorylation of tau and its intracellular aggregation into oligomeric species over time maturing into NFTs (**19**) which are the features we observe in short- and long-term TBI survivors, respectively.

Further research including a larger cohort of TBI cases, ranging from minutes or hours to years survival post-injury, is needed to validate our novel finding of tau oligomers formation following a head injury and to better characterise the appearance of beta amyloid plaques and tau pathology as well as the potential influence of genetic risk factors. In parallel, cellular and animal studies using tau oligomers isolated from human brain tissue (either post-mortem fresh-frozen tissue or biopsies from TBI patients) will be essential to determine whether these tau species are biologically active and capable of propagating pathology across the brain in a prion-like manner.

With TBI being a significant environmental risk factor for AD and reports that over 70 million people worldwide may be living with dementia by 2030, and the figure will double by 2050, is extremely important to understand the mechanisms leading from TBI to AD. Our findings of tau oligomers forming within 24 hours of head injury suggest that the sequence of the pathological events leading to tau pathology may be initiated decades before AD-like clinical symptoms emerge and could be amenable to therapeutic intervention. Notably, beyond serving as an excellent model for studying early mechanisms of AD, TBI also offers a unique opportunity for early therapeutic intervention in the disease process. In many cases, patients undergo neurosurgical procedures shortly after injury, offering rare access to the brain during a critical therapeutic window (e.g. microdialysis). If TBI initiates the prion-like spread of pathogenic tau oligomers, targeted therapies administered acutely could interrupt this cascade and potentially prevent the long-term development of AD. Notably, immunotherapies capable of blocking tau oligomer uptake are already in Phase 1 clinical trials (**20**). Thus, the identification of tau oligomers in acute TBI not only provides insight into early disease mechanisms but also highlights a promising therapeutic avenue for preventing AD.

## Method

### Ethic Statement

The study was undertaken using the ethics NRES 10/HO308/56 covering the use of anonymous samples of post-mortem human brain tissue.

### Subjects

The Cambridge Brain Bank provided anonymous post-mortem brain samples (paraffin embedded brain tissues) from TBI and AD cases as well as uninjured control cases known not to have any neurological or psychiatric disorder (**Table 1**). Different brain regions were obtained with the frontal, temporal and hippocampal regions available on all cases (**Supplementary Table 1**). Clinical data was retrospectively obtained from the clinical charts available at the Brain Bank and is summarised in Table 1. The healthy controls were assessed, by a qualified neuropathologist to be free of any Alzheimer pathology according to tau pathology and β-amyloid deposition (**21**).

### Immunohistochemistry

Immunohistochemistry was performed on 10μm thick paraffin embedded sections from TBI, AD as well as uninjured control brains using antibodies specific for NFTs, oligomeric tau and anti-amyloid plaques (**Supplementary Table 2**), following standard protocols (**6**). In particular, 10 brain sections from 11 brain regions were analysed (**Supplementary table 1**). Following deparaffinization and rehydration to dH_2_O, sections were immersed in 3% aqueous H_2_O_2_ to quench endogenous peroxidase activity. Then, following antigen retrieval via microwave and blocking using 1% normal horse serum, sections were incubated overnight at 4°C with the primary antibodies listed in Supplementary Table 2. The labelling was visualized the day after, with the ABC Elite Vectastain Kit (Vector laboratories). Briefly, sections were incubated for 2 hrs at room temperature with the biotinylated secondary antibody (1:500) and, following washes in PBS, horseradish peroxidase Avidin-D was added for 1 hr at room temperature and visualized with 3-3’diaminobenzidine as the chromogen. Controls omitting the primary reagent were included in all the experiments and were consistently negative for any staining.

### Semi-quantitative analysis

Semi-quantitative assessments of abnormal tau aggregates were conducted blind to the demographic and clinical information for all cases. Only AT8 (+) and T22 (+) tau positive profiles that could be clearly identified based on morphological characteristics (tangles and oligomeric tau) were counted in each section following previously published methods (**22, 23**). The average number of each tau profile (tau score) were determined across sections from the three brain regions (frontal, temporal and hippocampal) available for all cases and was categorised by applying the following ranked scale: ‘nearly absent’ (-, score 0), ‘sparse’ (+, score 1), ‘moderate’ (++, score 2), ‘high’ (+++, score 3), ‘very high’ (++++, score 4) and ‘extensive’ (>++++, score 5) (**Supplementary Table 3**). Tau pathology results were dichotomized in either ‘nearly absent/moderate’ score (**0–2**) or ‘high/extensive’ score (**3-5**) and the chi-square test was used to assess differences between groups (**4**).

## Acknowledgements

This research has received funding from Alzheimer’s Research UK (grant number ARUK-PPG2017B-020; RV and MC). The authors thank the Cambridge Brain Bank for the post-mortem tissue which is supported by a grant to the National Institute for Health Research (NIHR) Cambridge Biomedical Research Centre (KA). We are grateful to Robert Fincham and Olly Green at the Cambridge Brain Bank and Prof Nigel Temperton at the Medway School of Pharmacy for their technical assistance and support. Last but not least, our heartfelt gratitude to those donating their brain to support research.

## Author contributions

**S.K.B**. Performed immunohistochemistry, microscopy, statistical analysis and contributed to the writing of the first draft of the manuscript; **A.S**. contributed to the statistical analysis; **K.A**. contributed to the histopathology analysis; **M.C**. conceived and supervised the study; **R.V**. conceived and supervised the study, designed the research, performed the experiments, analysed the data and wrote the first draft of the manuscript. All the authors edited and proofread the manuscript.

