## Supplementary Figures and Tables for "Tau oligomers can occur in human brains within days of a single traumatic brain injury"

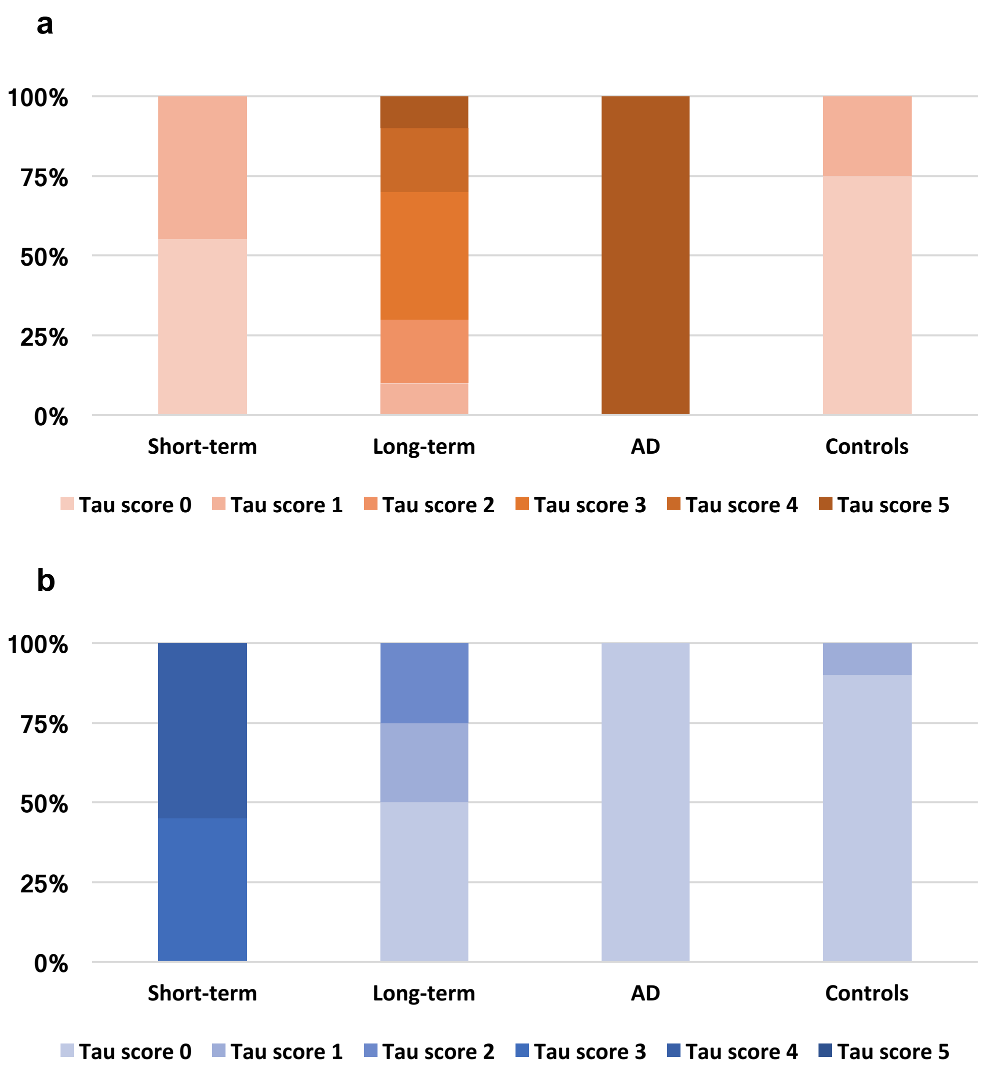


**Supplementary Figure 1 Combined semi-quantitative analysis of tau pathology and tau oligomers in short-term *versus* long-term TBI survivors.**
The analysis was determined on the three brain regions (frontal, temporal and hippocampal) available for each of the cases and categorised applying the following ranked scale: 'nearly absent' (score 0), 'sparse' (score 1), 'moderate' (score 2), 'high' (score 3), 'very high' (score 4) and 'extensive' (score 5). Results were dichotomized in either 'nearly absent/moderate' score (0 - 2) or 'high/extensive' score (3 - 5) and the chi-square test was used to assess differences between groups. (a) Tau pathology increases with time survival since the head injury. In particular, it is 'nearly absent/moderate' (score 0–2) in individuals who survived short-term whereas was 'high/extensive' (score 3–5) in long-term TBI survivors. (b) On the contrary, tau oligomers are 'high/extensive' (score 3-5) in short-term TBI survivors and 'nearly absent/moderate' (score 0-2) in long-term TBI survivors.

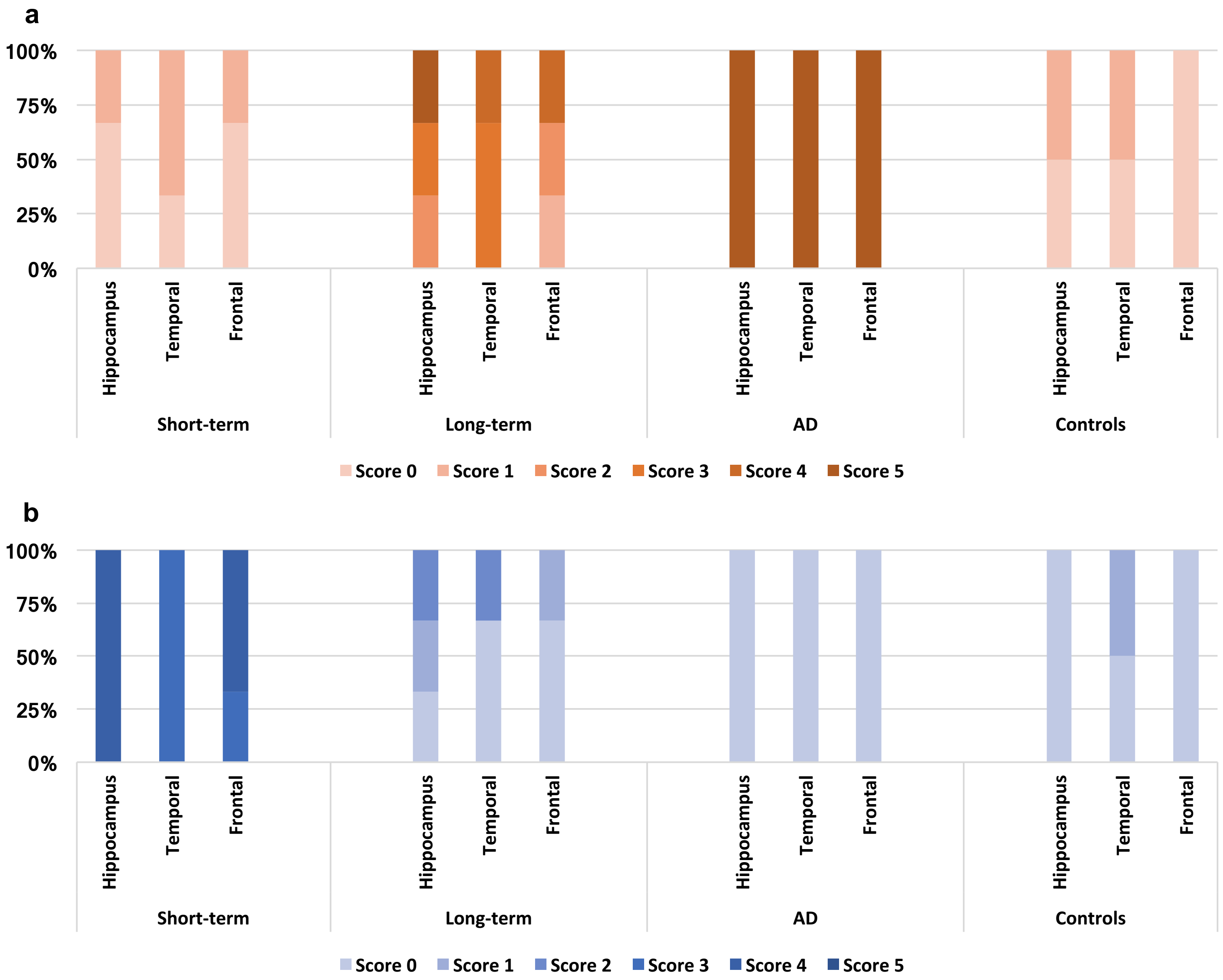


**Supplementary Figure 2 Semi-quantitative analysis of tau pathology and tau oligomers in the brain regions of short-term *versus* long-term TBI survivors**. The analysis was categorised applying the following ranked scale: 'nearly absent' (score 0), 'sparse' (score 1), 'moderate' (score 2), 'high' (score 3), 'very high' (score 4) and 'extensive' (score 5). Results were dichotomized in either 'nearly absent-moderate' score (0 - 2) or 'high-extensive' score (3 - 5) and the chi-square test was used to assess differences between groups. (a) Tau pathology increases with time survival since the head injury. In particular, it is 'high/extensive' (score 3-5)' in long-term survivors and 'nearly absent/moderate' (score 0 -2) in individuals who survived short-term. (b) On the contrary, tau oligomers are 'high/extensive' in short-term TBI survivors and 'nearly absent/moderate' in long-term survivors.

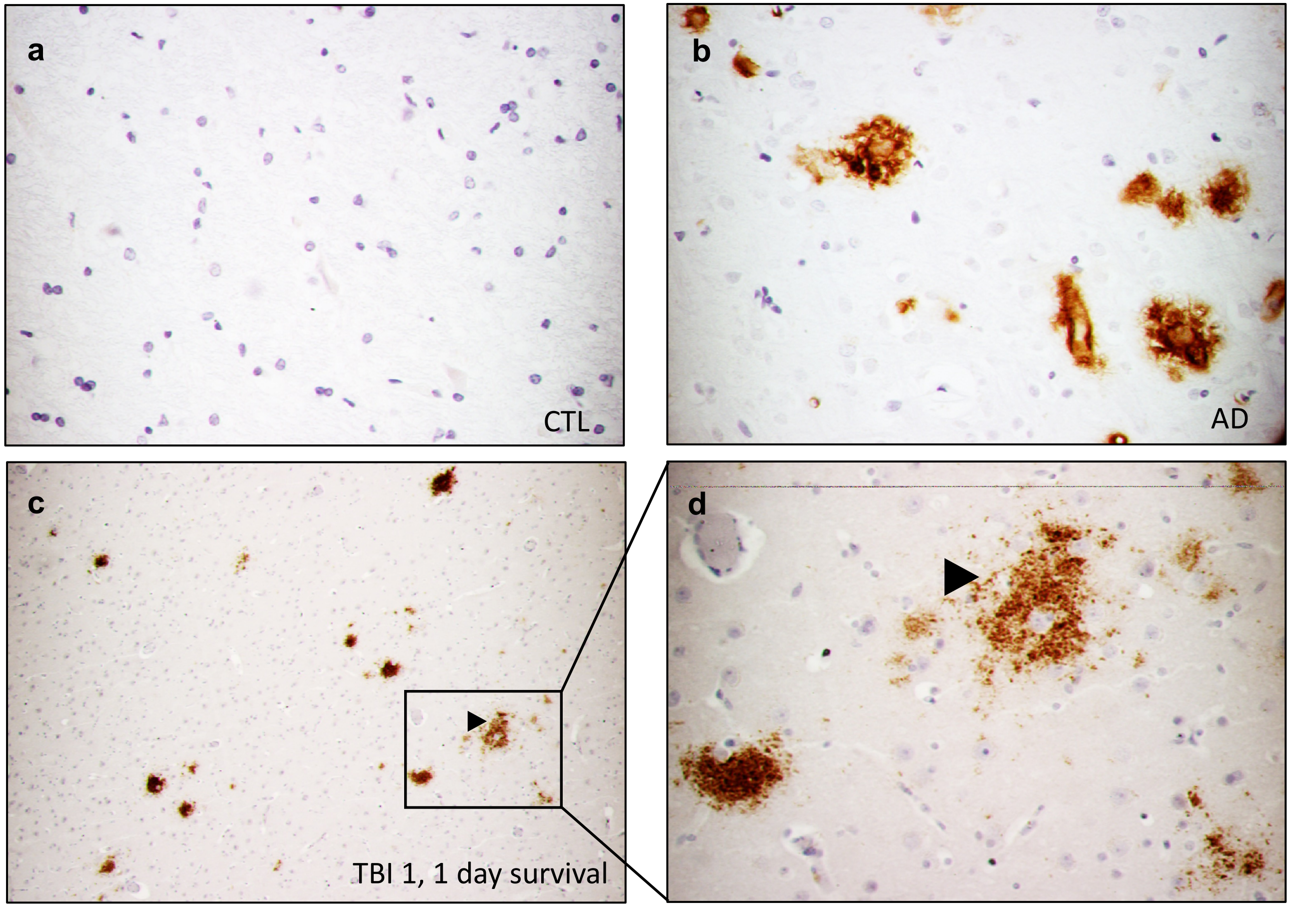


**Supplementary Figure 3 Amyloid Beta pathology in short-term TBI survivors.**
Immuno-histochemical detection of beta amyloid plaques in the frontal brain region of the individual aged 50 and with shortest time survival (TBI 1, 1 day, c-d) similar to those seen in AD (b). Amyloid plaques were absent in the uninjured control brain (a). Magnification 20X (c) 40X (a, b, d).

### Supplementary Table 1: Summary of brain regions

| Cases | Parietal | Corpus callosum | Cingulate | Hippocampus | Temporal | Frontal | Basal ganglia | Midbrain | Pons | Medulla | Cerebellum |
|---|---|---|---|---|---|---|---|---|---|---|---|
| **TBI 1** | | | | v | v | v | v | v | v | v | v |
| **TBI 2** | | | | v | v | v | | | v | | |
| **TBI 3** | | | | v | v | v | | | v | | v |
| **TBI 4** | | | | v | v | v | v | v | v | v | v |
| **TBI 5** | | | v | v | v | v | | | | | v |
| **TBI 6** | v | | | v | v | v | v | v | v | v | v |
| **AD** | | | | v | v | v | v | v | | | v |
| **CTL1**<br>**CTL2** | | | | v | v | v | v | v | v | v | v |

### Supplementary Table 2: Summary of antibodies used for Immunohistochemistry

| Antibody | Species | Dilution | Application | Epitope/Antigen | Source |
|---|---|---|---|---|---|
| **AT8** | Mouse monoclonal | 1:1000 | IHC | Tau (pSer202/Ser205) | Merck Millipore |
| **T22** | Rabbit polyclonal | 1:200 | IHC | Tau oligomers | Merck Millipore |
| **MOAB-2** | Mouse monoclonal | 1:1000 | IHC | Amyloid plaques | Merck Millipore |

**Supplementary Table 3: Semi-quantitative analysis of AT8 (+) and T22 (+) tau profiles**

| Cases (age) | Survival Time | Hippocampus | Temporal | Frontal |
|---|---|---|---|---|
| TBI 1 (50yo) | 1 day | ++++<br>- | +++<br>+ | ++++<br>- |
| TBI 2 (57yo) | 2 days | ++++<br>+ | +++<br>+ | ++++<br>+ |
| TBI 3 (26yo) | 2 days | ++++<br>- | +++<br>- | +++<br>- |
| TBI 4 (49yo) | 2 years | ++<br>++ | ++<br>+++ | +<br>+ |
| TBI 5 (58yo) | 6 years | +<br>+++ | -<br>+++ | -<br>++ |
| TBI 6 (67yo) | 30 years | -<br>+++++ | -<br>++++ | -<br>++++ |
| AD (78yo) | N/A | -<br>++++++ | -<br>++++++ | -<br>++++++ |
| CTL 1 (56yo) | N/A | -<br>+ | -<br>+ | -<br>- |
| CTL 2 (39yo) | N/A | -<br>- | +<br>- | -<br>- |
